# Unique actions of GABA arising from cytoplasmic chloride microdomains

**DOI:** 10.1101/2020.06.29.178160

**Authors:** Negah Rahmati, Kieran P. Normoyle, Joseph Glykys, Volodymyr I. Dzhala, Kyle P. Lillis, Kristopher T. Kahle, Rehan Raiyyani, Theju Jacob, Kevin J. Staley

## Abstract

Developmental, cellular, and subcellular variations in the direction of neuronal Cl^−^ currents elicited by GABA_A_ receptor activation have been frequently reported, and we found a corresponding variance in the reversal potential (E_GABA_) for individual interneurons synapsing on a single pyramidal cell. These findings suggest a corresponding variance in the cytoplasmic concentration of Cl^−^ ([Cl^−^_i_]). We determined [Cl^−^]_i_ by: 1) two-photon imaging of the Cl^−^ sensitive, ratiometric fluorescent protein SuperClomeleon (sCLM); 2) Fluorescence Lifetime IMaging (FLIM) of the Cl^−^ sensitive fluorophore MEQ; and 3) electrophysiological measurements of E_GABA_. These methods collectively demonstrated stable [Cl^−^]_i_ microdomains in individual neurons *in vivo*. Fluorometric and electrophysiological estimates of local [Cl^−^]_i_ were highly correlated. [Cl^−^]_i_ microdomains persisted after pharmacological inhibition of cation-chloride cotransporters (CCCs) but steadily decreased after inhibiting the polymerization of the anionic macromolecule actin. These studies highlight the existence of functionally significant neuronal Cl^−^ microdomains that modify the impact of GABAergic inputs.

## Introduction

The group-mean values of E_GABA_ are continuously distributed over a 25 mV range from −85 to - 60 mV in twenty recent studies utilizing gramicidin perforated patch recording techniques in CA1 pyramidal cells from rodents with mature neuronal Cl^−^ transport (Romo-Parra et al., 2008; Tyzio et al., 2008) **(Table S1)**. Over this wide range of reversal potentials, the effects of GABA_A_ receptor-gated currents will vary from inhibition via hyperpolarization of the membrane potential, through shunting inhibition, to excitation mediated by activation of low-threshold calcium currents and partial relief of the magnesium block of NMDA receptors (Doyon et al., 2016). These divergent effects suggest that E_GABA_ must be tightly regulated, although the mechanisms of regulation have not been resolved.

The observed range in E_GABA_ could be explained by a few mM variation in the cytoplasmic concentration of Cl^−^ ([Cl^−^]_i_), the anion with the highest permeability through GABA_A_Rs (Alfonsa et al., 2015b; Doyon et al., 2016; Raimondo et al., 2017). Such differences in [Cl^−^]_i_ could be maintained by the active transport of Cl^−^. For example, KCC2 and NKCC1 are oppositely-directed cation-Cl^−^ cotransporters (CCCs) expressed in neurons (Kahle et al., 2015). However, these are high-velocity transporters whose ionic equilibrium conditions do not match well with the distribution of observed E_GABA_ (Delpire and Staley, 2014), and inhibition of transport does not produce the predicted changes in [Cl^−^]_i_ (Glykys et al., 2014b; Sato et al., 2017).

Another possibility is suggested by the well-known partitioning effects on mobile ions exerted by the distribution of immobile ions (Fatin-Rouge et al., 2003) that for example create Gibbs-Donnan effects (Donnan, 1911) in gels (Procter, 1914) and unlinked biopolymers (Marinsky, 1985), and form the basis of ion exchange technologies (Yamamoto et al., 1988; Helfferich, 1995). Relatively immobile cytoplasmic biopolymers with high and spatially inhomogeneous densities of anionic charge, such as actin, tubulin, and nucleic acids (Gianazza and Righetti, 1980; Sanabria et al., 2006; Janke et al., 2008; Chen et al., 2020) could displace Cl^−^ locally (Glykys et al., 2014b; Glykys et al., 2017). Recent studies have underscored substantial variance in the subcellular distribution of these anionic biopolymers (Koleske, 2013; Morawski et al., 2015; Gut et al., 2018; Chen et al., 2020), suggesting that if the displacement of Cl^−^ underlies the intercellular diversity in E_GABA_, there should be a corresponding spatial variance in the subcellular distribution of Cl^−^. Indeed, many reports have suggested subcellular variance in [Cl^−^]_i_ or E_GABA_ (Pouzat and Marty, 1999; Berglund et al., 2006; Duebel et al., 2006; Szabadics et al., 2006; Romo-Parra et al., 2008; Földy et al., 2010; Glykys et al., 2014b; Astorga et al., 2015; Untiet et al., 2016; Zorrilla de San Martin et al., 2017; Schmidt et al., 2018) for which several mechanisms have been proposed (Khirug et al., 2008; Földy et al., 2010; Glykys et al., 2014b).

We report a unique E_GABA_ for each response to individual interneurons, and demonstrate a corresponding variance in local neuronal [Cl^−^]_i_ *in vivo* and *in vitro.* Several complimentary techniques were employed including two-photon microscopy of the ratiometric Cl^−^ indicator SuperClomelon (sCLM) (Grimley et al., 2013); Fluorescence Lifetime IMaging (FLIM) of Cl^−^- sensitive, pH-insensitive dye 6-methoxy-N-ethylquinolinium (MEQ) (Biwersi and Verkman, 1991); and simultaneous direct measurement of the reversal potential of locally-activated GABA_A_-gated membrane currents. We assessed the range, stability, transport dependence, and influence of anionic macromolecules on the observed variance in local baseline [Cl^−^]_i_.

## Results

### Unique response of single pyramidal cells to individual inhibitory interneurons

We recorded single pyramidal cells using whole-cell patch clamping while selectively activating individual neighboring inhibitory interneurons **(Figure 1A)**. We observed unique, reproducible values of E_GABA_ for each interneuron-pyramidal cell pair **(Figure 1B)** whose distribution paralleled the broad range of previously reported E_GABA_ **(Figure 1C and Table S1)**. Our results demonstrate that individual interneurons have unique E_GABA_ on postsynaptic pyramidal cells (n ; 40 cells, 10 experiments, ~23 mV range in E_GABA_). The measured range may be an underestimate of the variance of synaptic E_GABA_ in these neurons, because individual interneurons have on average 10 synapses onto pyramidal cells (Maccaferri et al., 2000; Bezaire and Soltesz, 2013), each of which may have a different E_GABA_, and the somatic recording would reflect the combined currents from all synapses originating from the activated interneuron. Not all stimulated interneurons evoked GABAergic currents in the recorded pyramidal cells. Fifty six percent of the stimulated interneurons resulted in recordable currents.

**Figure 1:**
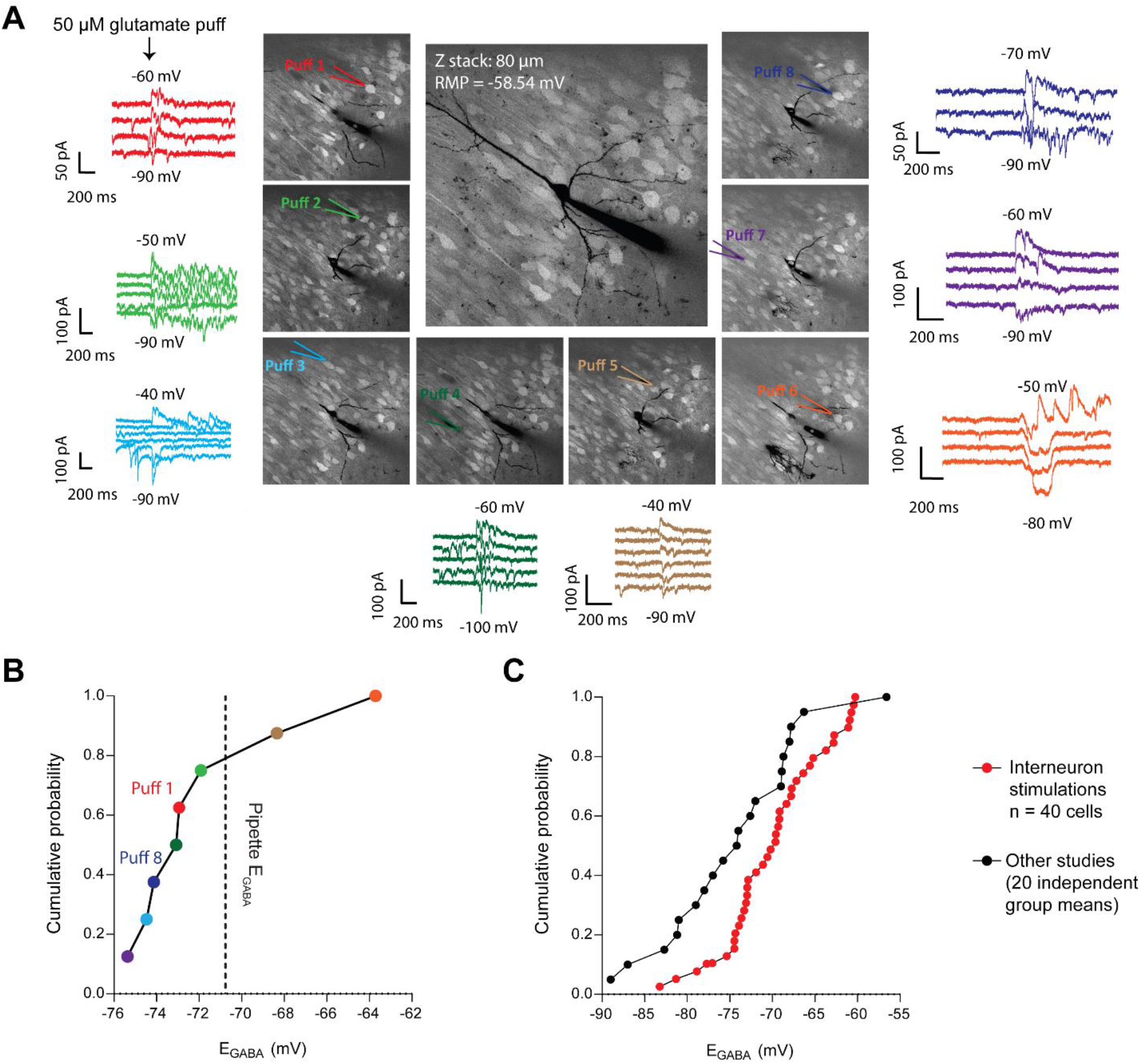
Inhomogeneity in responses to GABAergic interneurons. **A)** Puffing 50 μM glutamate to the soma of visualized interneurons (labeled with tdTomato) around a single pyramidal cell evokes GABA currents with different reversal potentials. **B)** Range of E_GABA_ measured in panel A. The colors of data points in panel B match the colors of traces and puff pipettes in panel A. The black dashed line shows the calculated E_GABA_ based on the Cl^−^ concentration inside the recording pipette (pipette E_GABA_). The puffing pipette was manually returned to the same area by the end of the experiment (puff 1 and puff 8 in panel A). The difference between the E_GABA_ evoked by puff 1 and puff 8 was less than 2 mV while the time interval between these two puffs was approximately 50 minutes. Although puff 8 was applied later than puff 1, its evoked E_GABA_ was further away from the pipette E_GABA_ compared to puff 1. These findings argue against the possibility that the [Cl^−^] inside the recording pipette homogenizes the dendritic [Cl^−^]_i._ **C)** Summary of our experiments (n ; 40 cells) and 20 separate studies indicating a wide rage in E_GABA_ in age-matched CA1 pyramidal cells.

### Evidence for dendritic chloride microdomains by fluorescence microscopy and electrophysiology

How can a single neuron express so many different values of E_GABA_? To investigate this, we measured the subcellular [Cl^−^]_i_ distribution in single neurons. To enable cell-type specific analysis of subcellular cytoplasmic Cl^−^ concentrations, we designed a conditional SuperClomeleon (*sCLM*) over-expression mouse line by generating a floxed stop/*sCLM* knock-in at the *Rosa26* locus (**Methods, Figure S1**). sCLM is a recently-developed ratiometric Cl^−^ indicator with improved sensitivity compared to its predecessor Clomeleon (CLM) (Grimley et al., 2013). These fluorophores measure cytoplasmic Cl^−^ in many neurons and subcellular locations at once, without the perturbations of the intracellular milieu induced by whole-cell and perforated patch clamping. DNA Sanger sequencing confirmed the presence of the knocked-in *sCLM* allele (**Methods, Figure S1**). Both heterozygous and homozygous mice were viable and fertile and survived to adulthood without any abnormalities.

*In vivo* ratiometric two-photon fluorescence imaging was performed on anesthetized mice in which sCLM expression was driven by crosses with either *CamKII* or *Dlx* Cre mice to selectively express the fluorophore in principal cells or interneurons, respectively (see **Methods**). YFP/CFP values were compared in neurons of cortical layers II and III **(Figure 2A)**. We analyzed the dendrites that were located at the same imaging depth **(Figure 2B)** to exclude the impact of depth-dependent differential scattering of cyan vs yellow light by the brain tissue (Boffi et al., 2018). The data are presented as YFP/CFP ratios rather than [Cl^−^]_i_ because calibration of sCLM is not feasible *in vivo* (Arosio and Ratto, 2014; Boffi et al., 2018). This is a consequence of limited capacity to permeabilize the membrane of target neurons and to manipulate the extracellular Cl^−^ concentration (Krapf et al., 1988). However, our data demonstrate a clear variance in YFP/CFP ratios in dendrites and soma **(Figure 2A and 2B)**.

**Figure 2:**
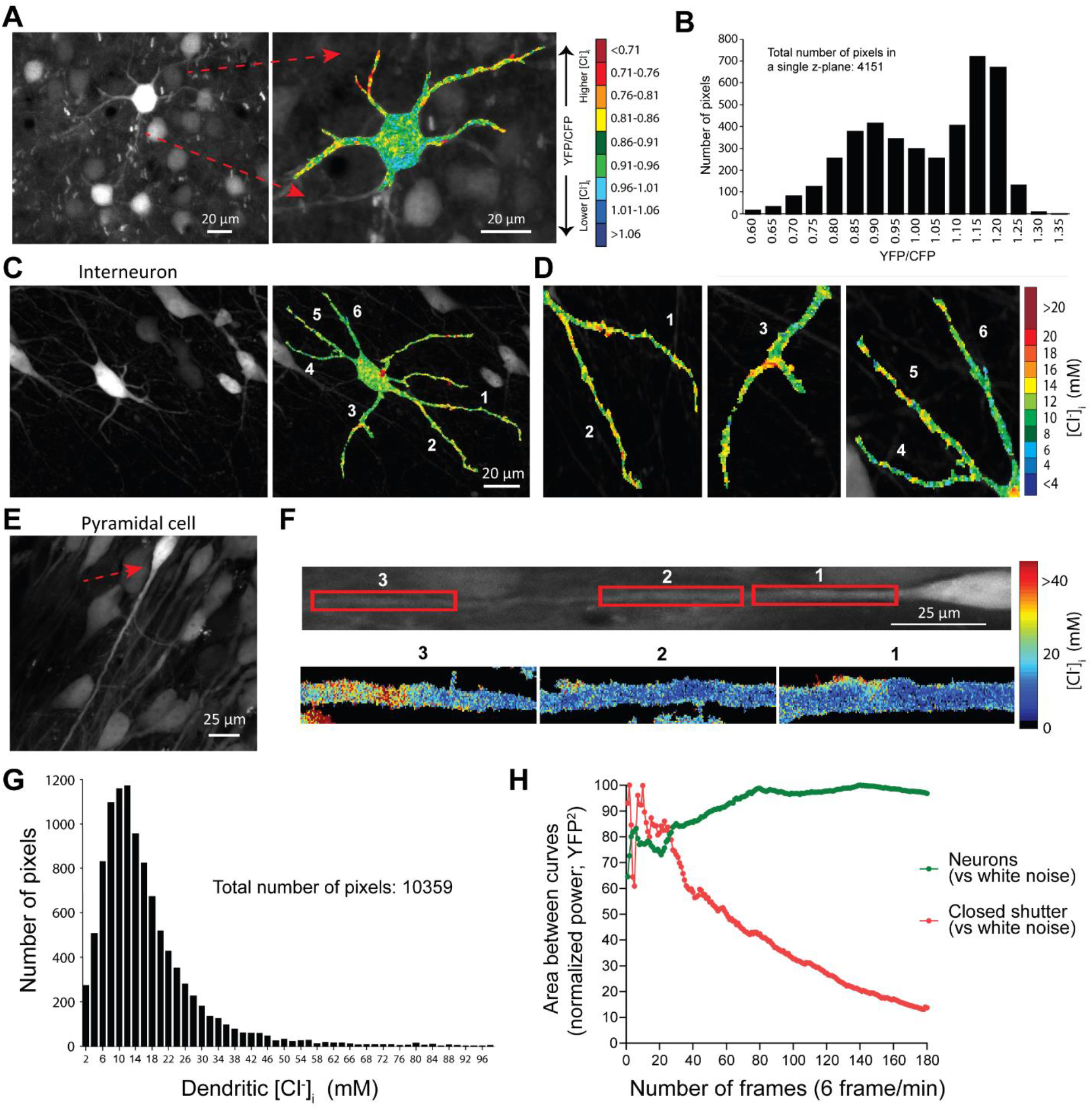
Evidence for the existence of chloride microdomains from sCLM imaging. **A)** A representative *in vivo* two-photon imaging of a layer III neuron expressing sCLM from a P20 mouse and the color-coded image of YFP/CFP. The image shows a 20 μm flatten z-stack. **B)** A histogram of somatic and dendritic YFP/CFP values restricted to a single z plane in the neuron shown in panel a. **C, D).** Two-photon imaging of an organotypic slice culture showing a hippocampal interneuron (C) and the dendritic variations in [Cl^−^]_i_ (D). **E)** A CA1 pyramidal cell from a hippocampal organotypic slice expressing sCLM. **F)** Top panel shows the rotated neuron in panel e with a red arrow. Bottom panel represents the magnified pseudo-colored [Cl^−^]_i_ maps of different segments of the dendrite. **G)** Histogram of all pixels derived from the dendrite of the neuron depicted in panel f. **H)** CLM expressing neurons were imaged for 30 minutes (180 frames at 6 frames per minute). The deviation from a white noise power spectrum is plotted against the number of timeframes averaged for the mean power spectra of both the neuronal YFP signal and closed-shutter noise. The resulting normalized curves converge toward significantly different values and demonstrate a decreased spatial variance over time in YFP signal vs normalized closed-shutter noise.

To analyze the subcellular Cl^−^ variance in a preparation in which sCLM YFP/CFP ratios can be quantitatively calibrated to [Cl^−^]_i_, we performed two-photon high resolution fluorescence microscopy of hippocampal organotypic slice cultures expressing sCLM. Our data demonstrate spatial variability of [Cl^−^]_i_ in the dendrites of individual neurons in the presence of 1 μM TTX **(Figure 2C-2F)**. The individual pixels of the neuron depicted in panel F demonstrate a wide range in dendritic [Cl^−^]_i_ **(Figure 1G)**. To examine the contribution of noise to these images, we performed time series imaging of CLM-expressing neurons (180 images over 30 minutes). Fluorescence intensity arising from stable cytoplasmic [Cl^−^]_i_ microdomains would be expected to exhibit less temporal variance than noise. Over the 30-minute observation period, the spatio-temporal variation in dendritic YFP intensity was much less than the corresponding variance produced by noise (measured with the shutter closed and normalized to the YFP emission intensity) **(Figure 2H and S2)**. This supports the presence of persistent spatial differences in [Cl^−^]_i_ that cannot be explained by noise.

sCLM retains the pH sensitivity of CLM (Grimley et al., 2013). To test whether the fluorescence measures of [Cl^−^]_i_ correlated with electrophysiological assays of [Cl^−^]_i_, simultaneous measurements of local dendritic [Cl^−^]_i_ by sCLM and direct calculation of E_GABA_ by gramicidin-perforated patch-clamp recording and local pressure application of 10 μM GABA were carried out in cultured hippocampal neurons **(Figure 3A)**. Local YFP/CFP ratios and [Cl^−^]_i_ measured by sCLM were highly correlated with [Cl^−^]_i_ calculated by E_GABA_ **(Figure 3B and 3C)**. The correlation coefficient between the two measures was 0.65 (n ; 38 ROIs, 11 cells, p < 0.0001), consistent with the role of cytoplasmic microdomains in setting the local E_GABA_ (Delpire and Staley, 2014; Glykys et al., 2017).

**Figure 3:**
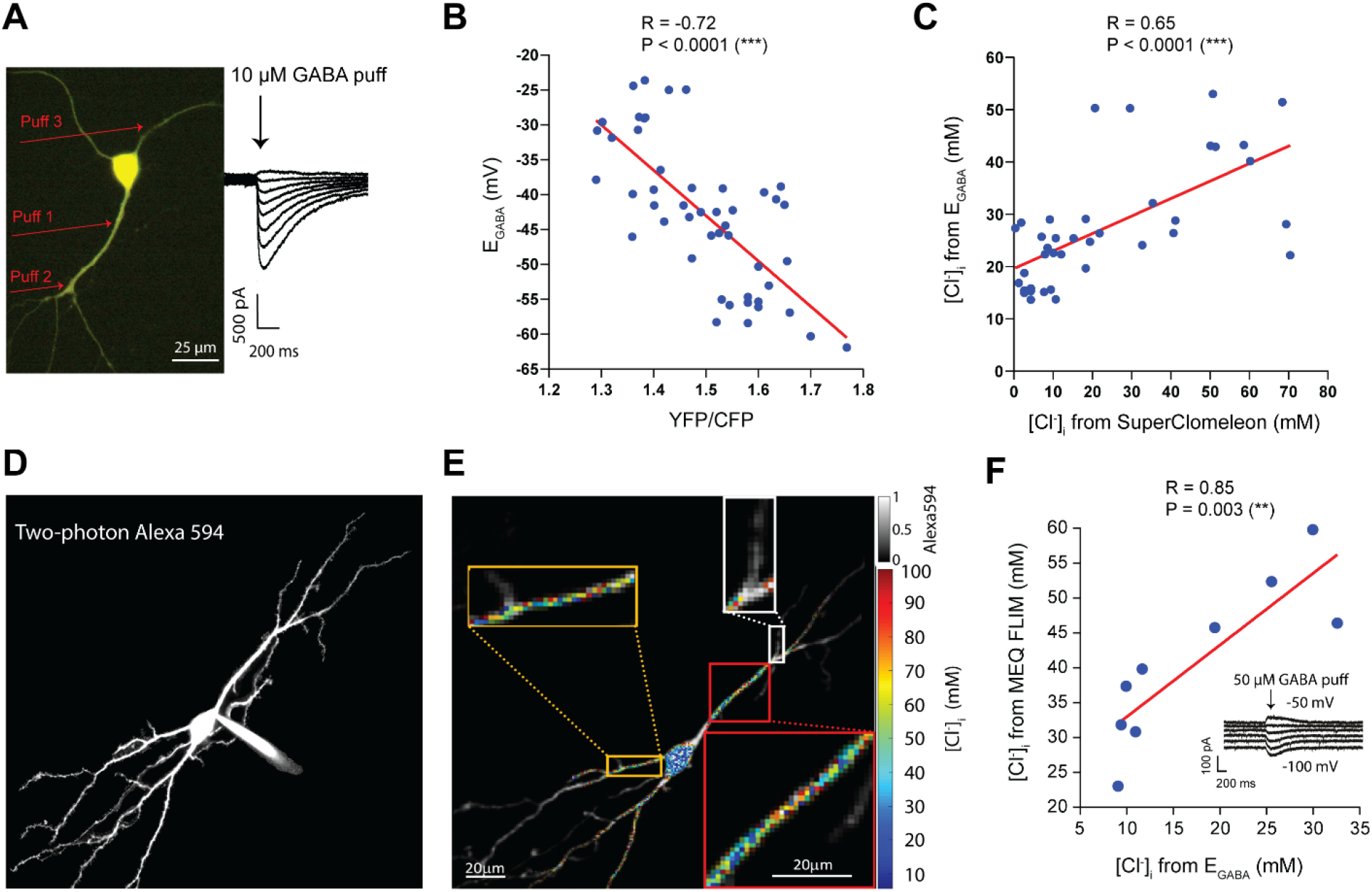
Simultaneous [Cl^−^]_i_ imaging and electrophysiology. **A)** A representative hippocampal dissociated cultured neuron expressing sCLM and visualized by DualView using a CCD camera to simultaneously measure the YFP and CFP fluorescence. **B)** The [Cl^−^]_i_ measured by fluorophore data (YFP/CFP) and the [Cl^−^]_i_ calculated from E_GABA_ in dissociated cultured cells are highly correlated (n ; 38 ROIs, 11 cells, p < 0.0001). **C)** There is a significant correlation between the [Cl^−^]_i_ measurements by sCLM and [Cl^−^]_i_ calculated by E_GABA_ (n ; 38 ROIs, 11 cells, p < 0.0001). **D)** A representative example of a hippocampal pyramidal cell loaded with Alexa Fluor-594 and Cl^−^-sensitive, pH-insensitive dye, MEQ. **E)** Fluorescence lifetime imaging of [Cl^−^]_i_ in hippocampal neurons confirms the variance of [Cl^−^]_i_ in dendrites. [Cl^−^]_i_ calculated from MEQ lifetime is visualized in pseudo color where the Alexa Fluor-594 ([Cl^−^]_i_-insensitive) signal sets the transparency value such that the [Cl^−^]_i_ value and corresponding color relate solely to the MEQ lifetime while the brightness of the pixel is set by Alexa Fluor-594 as a structural marker (gray colorbar, top right). No averaging or clustering was performed on MEQ images. Note the variety of [Cl^−^]_i_ values and the frequent juxtaposition of relatively high and low values in close proximity. **F)** After loading the cells with MEQ, 50 μM GABA was puffed to different visualized segments of dendrites and E_GABA_ was measured. The [Cl^−^]_i_ measured by MEQ lifetime and [Cl^−^]_i_ measured by E_GABA_ are significantly correlated (n ; 9 ROIs, 7 cells, p ; 0.003).

Utilizing another complimentary approach, we assayed subcellular [Cl^−^]_i_ using the pH-insensitive, Cl^−^ sensitive, single-wavelength fluorophore MEQ (Biwersi and Verkman, 1991). We performed FLIM to avoid artifacts arising from subcellular variation in MEQ concentration. The FLIM measurements using MEQ delivered by the whole-cell recording pipette confirmed the variability of [Cl^−^]_i_ in dendrites of hippocampal pyramidal neurons in CA1 area **(Figure 3D and 3E).** The variance in [Cl^−^]_i_ measured by MEQ FLIM was also highly correlated with simultaneous electrophysiological measurements of E_GABA_ measured in voltage clamp neurons by local application of GABA using direct visual guidance (n ; 9 ROIs, 7 cells; p ; 0.003) **(Figure 3F)**. These data support the idea that subcellular variance in [Cl^−^]_i_ is a significant contributor to the variance in E_GABA_ **(Table S1)**.

### Physiological chloride microdomains are stable and unaffected by pharmacological inhibition of cation-chloride cotransporters

Changes in [Cl^−^]_i_ are readily observed in response to alterations of the equilibrium conditions for CCCs using either osmotic or ionic perturbations of the perfusate (Thompson et al., 1988; DeFazio et al., 2000; Dzhala et al., 2010; Glykys et al., 2014b). This influence of transport equilibrium conditions on [Cl^−^]_i_ has led to the idea that CCCs set [Cl^−^]_i_ independently of the equilibrium conditions. To test the degree to which [Cl^−^]_i_ microdomains are defined by CCCs, we imaged sCLM in stable extracellular ionic and osmotic conditions. We acquired time-lapse images for 50-60 minutes **(Figure 4A)**. The ROIs (soma and dendrites of a single pyramidal neuron) were monitored and compared pixel by pixel with previous measurements every 10 minutes. Consistent with the prior noise analyses **(Figure 2H)**, the microdomains remained highly stable in control conditions. Applying a high concentration of furosemide to block NKCC1 and KCC2 simultaneously (Gillen et al., 1996) did not alter the microdomain distributions **(Figure 4A, 4C)**. Because the mobility of Cl^−^ in neuronal cytoplasm is very high (Kuner and Augustine, 2000), [Cl^−^]_i_ should have rapidly homogenized after blockade of CCCs if CCC activity had created the [Cl^−^]_i_ microdomains. The fact that this was not observed indicates that [Cl^−^]_i_ microdomains do not arise from local differences in cation-Cl^−^ transport rates. While the correlation coefficient was stable over one hour, the value of the correlation coefficient was dependent on the expression levels of sCLM in different mice **(Figure S5, panel D)**. This supports the idea that the spatial variance in sCLM signal is distinct from noise. To assess the lower bound of spatial stability that could be resolved with this technique, the same region of interest was acquired with the microscope shutter closed to quantify the much lower correlation of noise between the images **(Figure 4B and 4C)**. As a positive control to demonstrate that changes in dendritic [Cl^−^]_i_ could be measured with these techniques, dendritic [Cl^−^]_i_ was also measured in sCLM organotypic slices before and after seizure-like-events (SLE) (Glykys et al., 2014b). Our data show that the dendritic [Cl^−^]_i_ in 4 different ROIs return to the same baseline value after SLE **(Figure 4D and 4E)**, and that the return was independent of the initial (baseline) value. These results indicate that temporal fluctuations in [Cl^−^]_i_ can be resolved by this technique, and that [Cl^−^]_i_ microdomains persist after large transient Cl^−^ fluxes.

**Figure 4:**
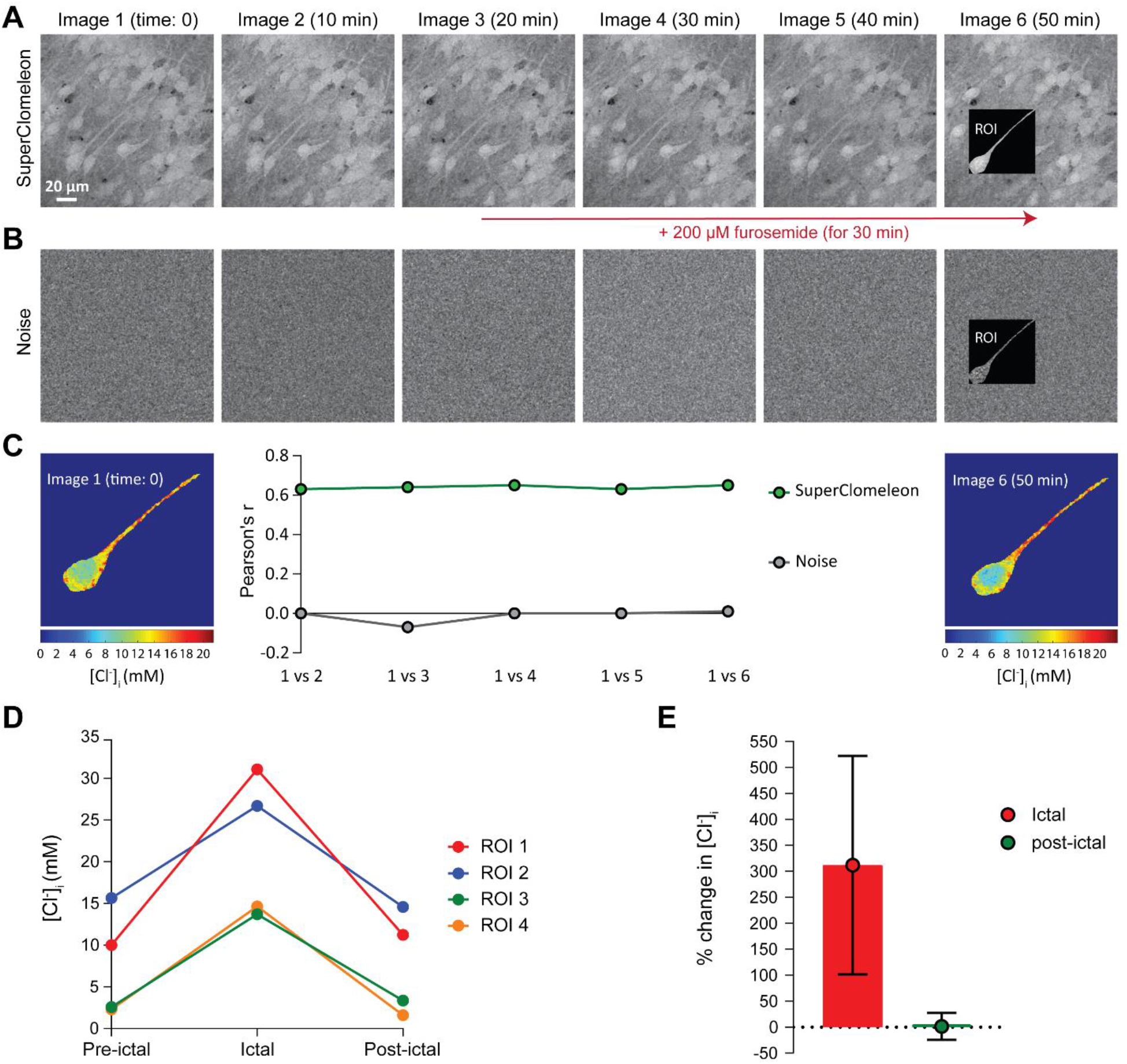
Chloride microdomains are stable and are not defined by Cl^−^ transport. **A)** Time-lapse images of a hippocampal pyramidal cell, expressing sCLM in the presence of 1 μM TTX before and after blocking KCC2 and NKCC1 with furosemide. **B)** The same region of interest was imaged while the microscope shutter was closed to measure the noise level and pixel by pixel correlation of the ROI as a control. **C)** The colocalization correlation analysis of the [Cl^−^]_i_ indicates that these dendritic Cl^−^ microdomains remain stable before and after the application of 200 μM furosemide. **D)**[Cl^−^]_i_ of four dendritic ROIs were measured before and after a spontaneous seizure in a sCLM organotypic slice. [Cl^−^]_i_ returns to the same baseline after the ictal period. **E)** Mean percent change in [Cl^−^]_i_ measured in the 4 ROIs shown in panel D. Data are presented as mean ± SD.

### The stability of chloride microdomains is reduced by actin depolymerization

If the observed cytoplasmic [Cl^−^]_i_ microdomains were created by the differential distribution of less mobile anionic macromolecules, then altering the distribution of those macromolecules should induce a corresponding change in chloride microdomains. To test this, we repeated the time-lapse two-photon imaging of sCLM organotypic slices (Figure 4) in the presence of latrunculin B which promotes net actin depolymerization (Morton, 2000) without affecting the morphology of dendrites (Kim and Lisman, 1999). To exclude secondary effects of cytoskeletal alterations on synapses and transporters, the experiments were performed in TTX, bumetanide (NKCC1 blocker) and VU 0240551 (KCC2 blocker). Following baseline imaging, neurons were continuously imaged for an additional hour after latrunculin B was added to the perfusate. Pixel by pixel correlation of dendritic and somatic ROIs from sequential images were compared with previous measurements **(Figure 5A)**. The application of latrunculin B steadily changed the distribution of microdomains, evidenced by a reduction in the pixel-wise correlation coefficient of sequential images **(Figure 5B)**. Over the same time period (40-60 minutes), the average percent change in pixel by pixel correlation in the presence of transporter blockers (n ; 12 ROIs, mean ± SD ; −1.09% ± 9.6%) was significantly lower, i.e. microdomain stability was higher, than in the presence of transporter blockers and latrunculin B (**Figure 5C**, n ; 13 ROIs, mean ± SD ; −19.45% ± 16.05%; p ; 0.002). The average pixel by pixel correlation was also compared between CFP images as control which did not significantly change during the same time period (mean transporter blockers ± SD ; −7.20% ± 15.5%; mean latrunculin B ± SD ; −7.09% ± 4.7%, p ; 0.98). The addition of latrunculin B to the perfusate containing transporter blockers did not alter the average percent change of [Cl^−^]_i_ observed after transport block **(Figure 5D**, mean transporter blockers ± SD ; 14.14% ± 24.2%; mean latrunculin B ± SD ; 14.74% ± 14.07%, p ; 0.94). This indicates that although CCCs are actively transporting Cl^−^ across the membrane in this preparation, the CCCs do not define the distribution of microdomains. In contrast, latrunculin B modified the distribution of [Cl^−^]_i_ micodomains without additional change to the average [Cl^−^]_i_ by altering the macromolecular distribution through biopolymer disassembly.

**Figure 5:**
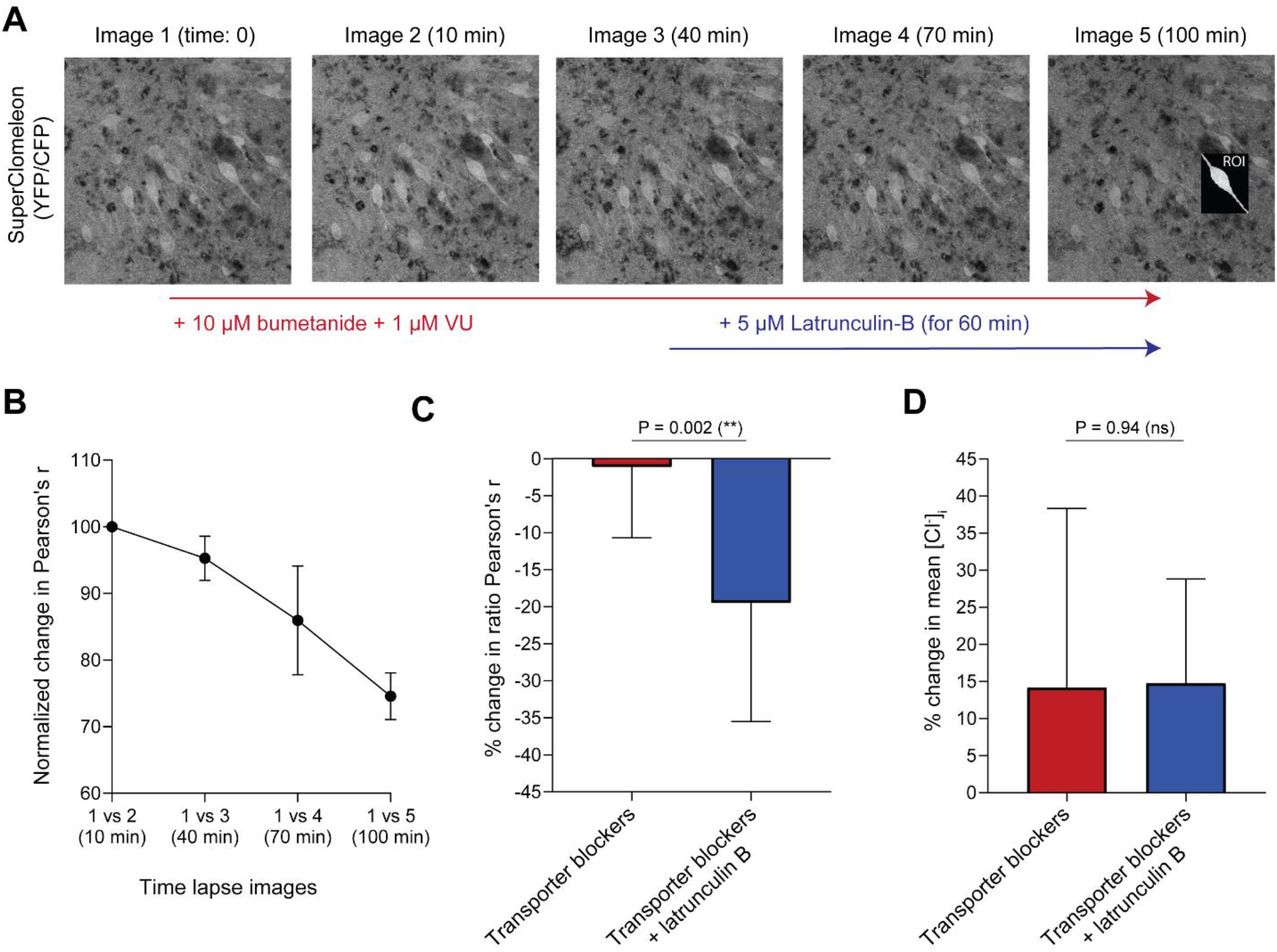
Contribution of cytoplasmic anionic immobile polymers to Cl^−^ microdomains. **A)** Time-lapse imaging of CA1 hippocampal sCLM expressing cells was performed in the presence of 1 μM TTX, 10 μM bumetanide and 1 μM VU. Latrunculin B was added to the perfusate and a pixel by pixel correlation was performed before vs. after drug-induced actin depolymerization. **B**) The change in colocalization correlation of the ratiometric images demonstrates that the spatial stability of Cl^−^ microdomains decreases after addition of 5 μM latrunculin B. **C)** The average pixelwise spatial correlation coefficient is significantly decreased after application of latrunculin B in addition to transport blockers compared to the time points when there are only transporter blockers in the bath. **D)**[Cl^−^]_i_ averaged over the entire ROI is not changed by the addition of latrunculin B to the perfusate. Data are presented as mean ± SD.

## Discussion

These data obtained using multiple complimentary recording techniques support the existence of stable [Cl^−^]_i_ microdomains and correspondingly unique responses to local GABA_A_ receptor activation in the dendritic and somatic cytoplasm of CA1 pyramidal cells. The stability of microdomains over the entire time interval for which stability could be assessed, and the wide variance in the E_GABA_ of pyramidal cells measured when individual presynaptic interneurons were stimulated, underscore the potential functional importance of [Cl^−^]_i_ microdomains for signaling at GABAergic synapses.

Neuronal Cl^−^ microdomains have been a source of confusion and controversy. A wide variety of electrophysiological evidence supporting the nonuniform distribution of cytoplasmic Cl^−^ (Barna et al., 2001; Duebel et al., 2006; Szabadics et al., 2006; Földy et al., 2010; Glykys et al., 2014b; Berglund et al., 2016; Untiet et al., 2016; Zorrilla de San Martin et al., 2017) has been challenged primarily by theoretical arguments (Luhmann et al., 2014; Doyon et al., 2016; Savtchenko et al., 2017; Düsterwald et al., 2018). The simplest argument for the existence of neuronal cytoplasmic Cl^−^ microdomains is the fact that the vast majority of anions in the cytoplasm are relatively immobile. In neurons, the GABA_A_-permeant ions Cl^−^ and HCO_3_^−^ together make up only a minority of the intracellular anions. The rest of intracellular anions are amino acids and phosphates (Morawski et al., 2015), of which only a small minority are not associated with macromolecules (Masuda et al., 1990; Veech et al., 2002) such as nucleic acid species (Manning, 1978) and proteins (Gianazza and Righetti, 1980). These immobile anionic polymers are not uniformly distributed in the cytoplasm (Gut et al., 2018; Chen et al., 2020), so it is reasonable to expect that there is a nonuniform distribution of mobile anions to compensate for the distribution of immobile anions. Macroscopic nonuniform distributions of mobile ions are readily demonstrated experimentally using gels with fixed anionic charges (Procter, 1914; Fatin-Rouge et al., 2003; Golmohamadi et al., 2012), and these findings extend to anionic biopolymers that are not crosslinked into gels (Marinsky, 1985). Here we demonstrate that inhibition of polymerization of one such anionic macromolecule, actin, led to a proportionate change in the distribution of Cl^−^ microdomains (Figure 5).

Cl^−^ is not the only cytoplasmic ion whose spatial distribution would be affected by immobile anionic polymers. Due to electrostatic interactions, the concentration of cations is much higher in the hydration shells of these anionic polymers than in the free water that surrounds the shell (Gregor, 1951; Marinsky, 1993). This cytoplasmic inhomogeneity in cation concentration may provide insight into a long-standing problem: how can the high-velocity, membrane CCCs be at equilibrium over the wide range of observed steady-state values of [Cl^−^]_i_ (Table S1; Figure 1C)? The simple ionic stoichiometry of the transporters (Voipio and Kaila, 2000), coupled with their high transport velocity (Thompson et al., 1988; Staley and Proctor, 1999; Jin et al., 2005; Alfonsa et al., 2015a) and presumed homogenous cytoplasmic cation concentrations should lead to homogenous [Cl^−^]_i_, but this is not observed (e.g. Figure 2; 1C; Table S1). However, if the local cytoplasmic cation concentration also varies, then a much wider range of local steady-state [Cl^−^]_i_ could be in equilibrium with the CCCs: cytoplasmic Cl^−^ microdomains (Figure 6).

**Figure 6:**
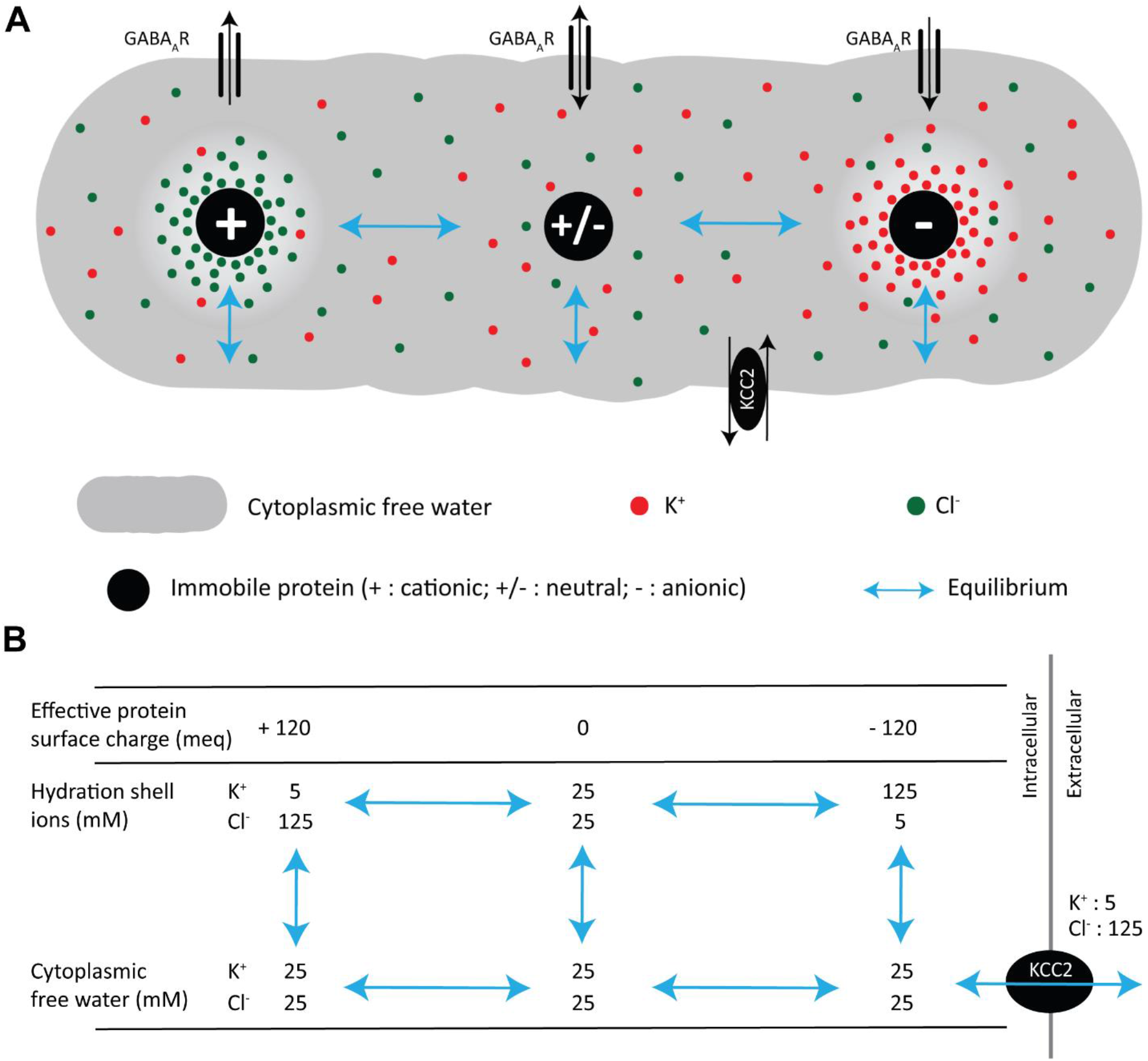
An example of the effects of nonuniform distribution of immobile anionic charge on the distributions of mobile cationic and anionic charges. **A)** All ions in the illustrated regions of the protein hydration shell are in equilibrium with the ions in the cytoplasmic free water. The Donnan criteria (local charge balance, and equal free energies across equilibria) are fulfilled between each region and the free water, and between all elements of the free water. There is no net diffusion between regions of the hydration shell because of the effects of the fixed protein surface charge. Most cytoplasmic proteins bear a substantial net negative surface charge, so the majority of cytoplasmic K^+^ resides in hydration shells, and the mean cytoplasmic Cl^−^ is less than the Cl^−^ in the free water. KCC2 is shown in equilibrium with the ions in the cytoplasmic free water. The additional osmotic pressure in hydration shells near high fixed surface charge density of the protein is a consequence of the locally higher concentration of counter-charged mobile ions and does not drive additional water movement (Helfferich, 1995), chapter 5). **B)** The concentrations shown in panel B are for illustration only; we do not yet know the actual ionic activities in the protein hydration shells or cytoplasmic free water. The example values for [Cl^−^]_i_ demonstrate that E_GABA_ can vary from −1 to −86 mV while maintaining equilibrium with KCC2-mediated KCl membrane transport.

The local variance of the concentrations (or more accurately, the activities) of local cations and anions provides a mean to reconcile the findings of transport biology (Voipio and Kaila, 2000; Doyon et al., 2016) with the impact of Donnan effects (Glykys et al., 2014b). That is to say, if transport is at equilibrium in each microdomain, then the highly regulated expression, phosphorylation and membrane trafficking of these transporters (Stein et al., 2004; Lee et al., 2010; Heubl et al., 2017; Garand et al., 2019) may serve to tailor transport to the characteristics of each microdomain, maintaining volume and average Cl^−^ flux.

A related question is: if impermeant anions rather than transporters set E_GABA_, why does inhibition of specific cotransporters cause small fractional changes in measured [Cl^−^]_i_ (Figure 5D) or E_GABA_ (Hübner et al., 2001; Dzhala et al., 2005; Sipilä et al., 2009)? First, E_GABA_ does not shift to resting membrane potential (Hübner et al., 2001; Brumback and Staley, 2008; Rinke et al., 2010), and [Cl^−^]_i_ does not shift to a passive distribution (Glykys et al., 2014b; Sato et al., 2017), as would be predicted if transporters set E_GABA_. Second, neuronal membrane permeability to Cl^−^ is comprised of multiple Cl^−^ cotransporters besides KCC2 and NKCC1, as well as GABA_A_ and glycine receptors, whereas in the system described by Donnan, the membrane ionic permeabilities are independent of each other (Donnan, 1911). In neurons, for each membrane Cl^−^ cotransporter, the Cl^−^ permeability is linked to transport of another molecular species with its own transmembrane free energy gradient. The sum of cotransport approximates an independent Cl^−^ permeability (Glykys et al., 2014b; Glykys et al., 2017), but pharmacological block of any one cotransporter will shift the equilibrium [Cl^−^]_i_ accordingly. Fluorescence imaging of large numbers of neurons demonstrate that these shifts occur in both directions after inhibition of each transporter (Glykys et al., 2014b; Sato et al., 2017), as would be expected if neurons have unique transmembrane ionic gradients. Inhibition of both major CCCs at the same time led to very modest effects on [Cl^−^]_i_ microdomains (Figure 4C and 5C), consistent with prior findings (Glykys et al., 2014b; Sato et al., 2017), whereas inhibition of actin polymerization significantly altered cytoplasmic Cl^−^ microdomains (Figure 5B and 5C).

The heterogeneity of [Cl^−^]_i_ provides a potential explanation for the higher [Cl^−^]_i_ reported by fluorophores vs. the [Cl^−^]_i_ recorded by electrophysiological measurements (e.g. Glykys et al., 2009; Dzhala et al., 2012; Sato et al., 2017; vs. Figure 1C). For example, a fluorophore sensing the Cl^−^ in cytoplasmic free water but not the Cl^−^ in hydration shells will return a value higher than the mean cytoplasmic Cl^−^. And if the fluorophore aligns with cationic regions of immobile proteins, it could report [Cl^−^]_i_ in those local hydration shells that are substantially higher than the mean cytoplasmic Cl^−^ (Figure 6).

Other proposed explanations for the heterogenous [Cl^−^]_i_ include variation in the stoichiometry of Cl^−^ cotransport (Brumback and Staley, 2008), obligatory water cotransport (Delpire and Staley, 2014; Glykys et al., 2019), and a low V_max_ of transport relative to membrane Cl^−^ flux (Doyon et al., 2016; Düsterwald et al., 2018). The last of these explanations appears to be a special case, because E_GABA_ at any synapse would be quite unstable within the physiological range of synaptic activity due to temporal fluctuations in synaptic Cl^−^ influx vs saturable cotransport at V_max_. Such temporal instability has only been observed at the maximal attainable levels of synaptic activity (Figure 4D, 4E; Huguenard and Alger, 1986; Staley et al., 1995; Köhling et al., 2000; Krishnan and Bazhenov, 2011; Alfonsa et al., 2015b; Silayeva et al., 2015; Buchin et al., 2016; Khazipov, 2016; Raimondo et al., 2017; Burman et al., 2019). The data presented here are most consistent with the idea that the equilibrium value of [Cl^−^]_i_ is defined by the local distribution of cytoplasmic anionic biopolymers. CCCs maintain that equilibrium value in the face of anionic membrane currents. This role for active Cl^−^ cotransport becomes critical as the rate of synaptic activity increases (Staley and Proctor, 1999; Glykys et al., 2009; Glykys et al., 2014b; Doyon et al., 2016). The proposal that the observed Cl^−^ distribution by displacement is only relevant at low rates of synaptic activity *in vitro* (Doyon et al., 2016; Düsterwald et al., 2018) is not consistent with *in vivo* measurements (Figure 2A, 2B). *In vitro* and in the presence of TTX, low rates of synaptic activity should dramatically reduce the discrepancy between measured E_GABA_ and the equilibrium conditions for cation-Cl^−^ cotransport if active transport were the primary mechanism of setting E_GABA_, but this has not been observed (Figure 2; Table S1; Glykys et al., 2014b; Sato et al., 2017).

The range of variance of E_GABA_ determined by activation of local interneurons agrees well with the ranges observed using perforated patch techniques (Figure 1). This raises the possibility of actively maintained motifs of immobile anions near the intra- and extracellular faces of the channels opened by GABA_A_ receptors. The range of E_GABA_ could be actively maintained by the local density and post-translational modifications of anionic intracellular structural species, such as gephyrin, actin, and tubulin (Sola et al., 2001), and extracellular sulfated glycosaminoglycans (Glykys et al., 2014a, b; Glykys et al., 2017).

The wide range in E_GABA_ directly impacts the direction and magnitude of the response to GABA released from each interneuron, and potentially at each GABAergic synapse. This increases the range of possible interactions between GABAergic and excitatory inputs to include not only hyperpolarizing and shunting inhibition but also synapse-specific amplification (Gulledge and Stuart, 2003; Grienberger et al., 2017). The observed 25 mV range in E_GABA_ is continuously distributed. This distribution is not consistent with a result arising from inadvertent stimulation of two interneuron subtypes with distinct E_GABA_ (Szabadics et al., 2006; Földy et al., 2010; Armstrong and Soltesz, 2012). Our findings do not invalidate those studies regarding E_GABA_ vs interneuron subtypes. Rather, interneuron subtype may be one of several important variables that determine E_GABA_ at each pyramidal cell GABA_A_ synapse.

Cl^−^ microdomains were stable over an hour, which was the longest we could feasibly measure. Future studies to elucidate the cellular and synaptic specificity of the polarity of GABA signaling, the time range over which this polarity is stable, and its plasticity will clarify the role of neuronal cytoplasmic Cl^−^ domains in neuronal signal processing and pathological states, such as medically intractable epilepsy (Cohen et al., 2002; Spruston et al., 2016).

## Acknowledgments

KS was supported by NIH/NINDS 5R01NS40109-14. JG was supported by NIH/NINDS 1K08NS091248.

## Author contributions

N.R. and K.J.S. designed research; N.R., K.P.N., J.G., and V.I.D. performed research; N.R., K.P.N., R.R. and T.J. analyzed data; N.R. and K.J.S. wrote the manuscript; K.P.N., J.G., K.P.L., V.I.D., K.T.K and R.R. participated in discussion and editing the manuscript.

## Declaration of interest

The authors declare no competing interests. Dr. Jacob is currently an employee of Bayer Corporation.

## Methods

### Ethics statement

All experiments were performed in accordance with protocols approved by the Center for comparative medicine (CCM) at Massachusetts General Hospital (MGH) and in accordance with the National Institute of Health Guide for the Care and Use of Laboratory Animals.

### Mice

#### Construction of the *sCLM* targeting vector

The STK1-HR gene-targeting vector was constructed from 129Sv mouse genomic DNA (GenOway, Lyon, France). The final targeting vector has the following features: (i) isogenic with 129Sv ES cells, favoring the homologous recombination; (ii) asymmetrical homology arms (5’ short arm-SA: 1 kb, 3’ long arm-LA: 4.3 kb); (iii) transgenic cassette containing the sCLM-pA elements allowing the expression of the transgene under the control of the ubiquitous CAAG promoter sequences; (iv) LoxP flanked, combined STOP-neomycin selection (Neo) cassette allowing its excision through the action of Cre-recombinase, activating the sCLM over-expression; (v) presence of the Diphtheria Toxin A (DTA) negative selection marker reduces the isolation of non-homologous recombined ES cell clones **(Figure S1A)**.

### Production of sCLM targeted ES cell clones

Linearized targeting vector was transfected into 129Sv embryonic stem (ES) cells (GenOway, Lyon, France) according to GenOway’s electroporation procedures. Polymerase chain reaction (PCR), Southern blot, and sequence analysis of G-418–resistant ES clones revealed the recombined locus in two clones. PCR across the 5′ end of the targeted locus used a forward primer hybridizing upstream of the 5′ homology arm (5′-AAGACGAAAAGGGCAAGCATCTTCC-3′) and a reverse primer hybridizing within the neomycin cassette (5′-GCAGTGAGAAGAGTACCACCAAGAGTCC-3′). Two Southern blot assays were hybridized with an internal and an external probe to assess recombination accuracy at the respective 5’ and 3’ ends of the *sCLM* locus. The absence of off-target mutations was confirmed by sequence analysis.

### Generation of chimeric mice and breeding scheme

Recombined ES cell clones microinjected into C57BL/6 blastocysts gave rise to male chimeras with statistically significant ES cell contribution. These chimeras were bred with C57BL/6J mice expressing Cre recombinase to produce the *sCLM* heterozygous line lacking the neomycin cassette. F1 genotyping was performed by PCR and Southern blot. PCR primers hybridizing upstream (5′-GGGCAACGTGCTGGTTATTGTGC-3′) and downstream (5′-ACAGCTCCTCGCCCTTGCTCAC-3′) of the neomycin cassette allowed PCR identification of the 3195–base pair (bp) *sCLM* allele amplicon harboring the neomycin cassette, and the 177-bp amplicon lacking the neomycin cassette. Southern blot hybridization with an external probe allowed identification of the 3.7-kb *sCLM* allele (**Figure S1B, C**).

Ratiometric [Cl^−^]_i_ measurements were performed on strain 129 sCLM mice (Grimley et al., 2013) by crossing sCLM-floxed mice with C57/BL6 Synapsin-Cre (for neuronal expression) or C57/BL6 DLX-Cre (for interneuron expression). FLIM experiments were performed on C57/BL6 wild type (WT) mice. Functional studies of [Cl^−^]_i_ microdomains were performed on DLX-Cre mice to visualize interneurons. Either sex was used for experiments.

### *In vivo* imaging

Craniectomies were done under aseptic conditions, under anesthesia with inhaled isoflurane and in a stereotaxic frame. The temperature was regulated through a heating pad. The surgical site was sterilized with betadine and isopropyl alcohol, and a 2–3 mm incision was made in the scalp along the midline between the ears. A 5-7 mm diameter hole was drilled in the skull, 1 mm lateral, and posterior to Bregma (parietal region) using a high-speed micro-drill. Once the craniotomy is completed, a thin round cover glass (~100 microns) is secured to the bone with a mixture of superglue and dental cement. Immediately after the craniectomy, the mice were imaged under the two-photon microscope while maintaining anesthesia and normo-temperature.

### Organotypic hippocampal slice preparation

P6-P7 mice were used for preparation of organotypic slice cultures using the rocking plate technique (Romijn et al., 1988) or the membrane insert technique (Stoppini et al., 1991) for both electrophysiology and imaging. Isolated hippocampi were cut into 400 μm slices on a McIlwain tissue chopper (Mickle Laboratory Engineering). Slices were then transferred to membrane inserts (PICMORG50; Millipore) or coverslips, which were placed in glass-bottomed six-well plates (P06-1.5H-N, CellVis). Both culture configurations were incubated at 5% CO2, 36°C in Neurobasal-A growth medium supplemented with 2% B27, 500 μM GlutaMAX, and 0.03 mg/ml gentamycin (all from Invitrogen). Growth medium was changed every 3-4 day. Slices were used for experiments between DIV 6 and DIV 20.

### Dissociated cell culture preparation

Hippocampal murine neurons were prepared based on the method of Beaudoin and colleagues (Beaudoin III et al., 2012), modified to use early post-natal mice and to employ a commercial kit (Worthington Biochemical, LK003150). P1-P4 mice are sacrificed by guillotine method cut just caudal to the ears. Bilateral incisions are made from ear to eye as far ventral as feasible, allowing rostral reflection of the calvaria. Using a spatula dipped in Gey’s balanced salt solution (GBSS; Sigma, G9779) supplemented with glucose (Sigma) to 1.5% and then filter sterilized, brains from up to four animals are gently removed and to a 6cm petri dish of glucose-supplemented GBSS in 6 cm petri dish on ice. Each hippocampus is dissected out and transferred to a second 6 cm petri dish of glucose-supplemented GBSS on a cold pack within the prep hood. Using a commercially available papain tissue dissociation kit (Worthington Biochemical, LK003150), hippocampi are transferred to Earle’s Balanced Salt Solution (EBSS)-based papain solution supplemented with DNase in a 6cm petri dish pre-equilibrated in 5% CO_2_ 36°C incubator and chopped for 30 seconds with Bonn scissors before being returned to incubator for a 10-to 40-minute incubation commiserate with pup age. Halfway through incubation, tissue is triturated x5 with a 10mL plastic pipette and returned to the incubator. Note that reduced yield from using plastic pipettes is deemed acceptable in order to avoid inconsistency of flame-polished glass tip diameter and subsequent trituration-related damage. Tissue is again triturated x5 with a 10 ml pipette and x3-4 with a 5 ml pipette before being strained through a 40-micron basket filter (Corning, 431750) and transferred to a 50 ml conical tube. Cell mixture is centrifuged 6 minutes at 300 x g and after carefully aspirating the supernatant the cell pellet is then resuspended with EBSS plus ovomucoid inhibitor albumin mixture supplemented with DNase (each part of Worthington kit). Gentle trituration with a 10 ml pipette held several centimeters from the tube bottom is sufficient to resuspend. This mixture is then carefully and slowly overlaid atop 5 ml of Ovoid protein in EBSS and centrifuged for 5 minutes at 70 x g. The resulting pellet is resuspended in Neurobasal-A media (Gibco, 10888-022) without supplementation and transferred to a fresh tube. Cell concentration and proportion surviving are calculated using trypan blue and a hemocytometer, and pre-plated to poly-D-lysine- and laminin-coated (Sigma, P0899 and laminin L2020, respectively) 19 mm coverslips (Electron Microscopy Services) or gridded glass-bottom 3.5 cm petri dishes (Ibidi, 81168) delivering 300 viable neurons per square millimeter at overall amounts between 2×10^5^ and 5×10^5^ cells in the absence of media supplementation. After a pre-plating period of two hours during which neurons will preferentially attach to the coated surface, unattached cells are aspirated, and fresh Neurobasal-A media supplemented as described above (Organotypic Hippocampal Slice Preparation). Fresh media replaces half of full volume every 2 to 3 days. Neurons were imaged between DIV 10 and DIV 20.

### Chloride measurements

We used three methods to measure [Cl^−^]_i_:

1. <u>Two-photon sCLM imaging</u>: The <u>sCLM</u> (Grimley et al., 2013) variant of CLM (Kuner and Augustine, 2000) was used for non-invasive [Cl^−^]_i_ imaging. <u>sCLM</u> consists of two fluorescent proteins CFP and YFP, joined by a short polypeptide linker which allows FRET-based imaging of [Cl^−^]_i_ (Grimley et al., 2013). For the Cl^−^ imaging experiments that were performed without electrophysiological recordings, two-photon imaging was performed using a Fluoview 1000MPE with prechirp optics and a fast acousto-optical modulator (AOM) mounted on an Olympus BX61W1 upright microscope equipped with a 25× 1.05 NA water-immersion objective (Olympus). A mode-locked Ti:Sapphire laser (MaiTai; Spectra-Physics, Fremont, CA) generated two-photon fluorescence with 860 nm excitation. Emitted light was detected through two filters in the range of 460-500 nm for cyan fluorescence protein (CFP) and 520-560 nm for yellow fluorescence protein (YFP). Two photomultiplier tubes (Hamamatsu Photonics) were used to simultaneously acquire CFP and YFP signals. Three dimensional stacks (3D) of raster scans in the XY plane (0.33 μm/pixel XY) were imaged at z-axis interval of 1 μm for measuring the stability of microdomains (Figure 4 and 5) and a z-axis interval of 2 μm for the rest of experiments. The pyramidal cell shown in Figure 1E was imaged with a higher resolution (0.099 μm/pixel XY and z-axis interval of 1 μm). Simultaneous two-photon and electrophysiological recordings were performed with a custom-built scanning microscope. Two-photon images were acquired using a custom-designed software (LabVIEW) and a scan head from Radiance 2000 MP (Bio-Rad), equipped with a 40×, 0.8 NA water-immersion objective (Olympus), and photomultiplier tubes with appropriate filters for CFP (450/80) and YFP (545/30). A Spectra-Physics Mai Tai laser was set to 860 nm for sCLM imaging. Serial images were collected with z-axis interval of 2 μm. The images were reconstructed offline either by ImageJ (PRID: SCR-003070) or MATLAB (v2017a). To calculate [Cl^−^]_i_ using Image J, a region of interest (ROI) was drawn around the cell body and dendrites and the ratio of YFP/CFP fluorescence intensity was measured. Images analyzed with MatLab were masked using image processing functions as described below. In either case the ratio was converted into [Cl^−^]_i_ by the following equation:

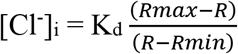

Where K_d_ is the dissociation constant, R_max_ is the ratio obtained in the absence of Cl^−^ and R_min_ is the ratio when <u>sCLM</u> is maximally quenched (Grimley et al., 2013) (Figure S3A).

2. <u>Fluorescent Lifetime IMaging (FLIM) of MEQ</u>: 1.5 mM MEQ (6-methoxy-N-ethylquinolinium iodide) was delivered to the cytoplasm via whole-cell recording pipette (Figure 2). Unlike <u>sCLM</u>, MEQ is insensitive to pH (Biwersi et al., 1992). The Cl^−^-sensitive signal of both fluorophores used in this study is independent of the concentration of dye due to either the ratiometric nature of sCLM, or the intrinsic nature of fluorescence lifetime as a concentration-independent property. MEQ was excited at 750 nm using the custom-built scanning microscope detailed above with appropriate emission filter (390/65) and fluorescent lifetime was measured using Becker & Hickl SPCImage software (Becker & Hickl, Berlin, Germany). The selection of the ROI and calculation of MEQ lifetime was measured offline by a custom-written MATLAB (MathWorks Inc. Natick, MA) program. MEQ lifetime values were translated to [Cl^−^]_i_ using Stern-Volmer calibration (Figure S3B) (Biwersi et al., 1992). The detailed morphologies of the cells were visualized by adding 20 μM Alexa Fluor-594 to the recording pipette. High-resolution Alexa Fluor-594 images (0.307 μm/pixel XY; 2 μm/pixel Z) were used for precise selection of the ROI in MEQ lifetime images with XY resolution of 1.537 μm/pixel and Z-axis interval of 2μm.

3. <u>Electrophysiological measurement of EGABA</u>: A subset of experiments were performed using whole-cell and gramicidin perforated patch-clamp techniques (details in the electrophysiology section) in conjunction with <u>sCLM</u> imaging or FLIM. The reversal potential of GABA_A_-current (E_GABA_) was extracted from currents evoked by puff application of GABA to different segments of the dendrites while clamping the cell at various membrane potentials. GABA_B_Rs were blocked by 2 μM CGP55845. The size of the dendritic region of interest was determined by adding Alexa Fluor-594 inside the puffing pipette and measuring the region that the dye was diffused. ([Cl^−^]_i_ was calculated using the Nernst equation:

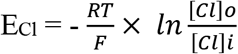

where R is the gas constant (8.315 J.mol^−1^. K^−1^), T is the temperature in Kelvin, F is the Faraday’s constant (96.487 C.mol^−1^), [Cl^−^]_i_ is the intracellular concentration of chloride and [Cl^−^]_o_ is the extracellular concentration of chloride.

### Electrophysiology

For electrophysiological recordings, organotypic slice cultures or dissociated cultured cells were transferred to a recording chamber and perfused with ACSF (2.5 ml/min) containing the following (in mM): 124 NaCl, 1.25 NaH_2_PO_4_, 2.5 KCl, 26 NaHCO_3_, 2 CaCl_2_, and 2 MgSO_4_ and 20 D-glucose, bubbled with 95% O_2_ and 5% CO_2_ at 34°C. Two separate series of electrophysiological experiments were performed: 1. Gramicidin perforated patch-clamping on hippocampal dissociated cell cultures: cells were visualized using an upright microscope (Nikon Eclipse FN1) equipped with a 40×, 0.8 NA water-immersion objective (Nikon), a DualView image intensifier (Optical Insights, LLC) and a CCD camera (Chameleon3, Point Grey Research Inc.) for simultaneous acquisition of two separate images of CFP and YFP with X resolution of 1.996 μm/pixel and Y resolution of 1.896 μm/pixel (Figure 3A). Electrodes were pulled from borosilicate glass capillaries (Sutter Instruments) using a micropipette puller (model P-97, Sutter Instruments) with resistance 5-7 MΩ when filled with internal solution containing the following (in mM): 140 KCl, 5 MgCl_2_, 10 HEPES, 5 EGTA, osmolarity 290 mOsm, pH 7.25-7.35, adjusted with KOH. Gramicidin-perforated patch clamping was performed as described previously (Rahmati et al., 2016). 2. Whole-cell patch-clamp recoding of hippocampal organotypic slices: These experiments were performed on wild type C57 rocking plate organotypic slices to simultaneously load the hippocampal pyramidal cells with MEQ for FLIM imaging and measure E_GABA_ at different segments of the dendrites by 50 μM GABA puff application (Figure 3E, F). Whole-cell patch-clamp recordings were also performed for functional evaluation of Cl^−^ microdomains (Figure 1) on organotypic slices of DLX-Cre mice which were transfected on the day of slicing with tdTomato virus (Addgene-AAV9-CAG-FLEX-tdTomato, 2 μl/ml culture media) to visualize interneurons. Patch pipettes with resistance 5-7 MΩ were filled with an internal solution containing the following (in mM): 124 K-MeSO_4_, 5 KCl, 10 KOH, 4 NaCl, 10 HEPES, 28.5 sucrose, 4 Na_2_ATP, 0.4 Na_3_GTP, 1.4 mM MEQ, osmolarity 295 mOsm, pH 7.25-7.35. For visualizing the whole morphology of the recorded cells, 20 μM Alexa Fluor-594 hydrazide (Invitrogen) was added to the internal solution on the day of the experiment. Resting membrane potential (RMP) and input resistance were measured after whole-cell configuration was reached. Series resistance (R_s_; assessed in voltage-clamp mode by −5 or −10 mV voltage step) was monitored during the experiments to be stable. Only neurons with R_s_ < 20 MΩ were included in the analysis. For all the electrophysiological recordings, signal acquisition was performed using a Multiclamp amplifier (Multiclamp 700B, Molecular Devices) with Clampex 10 software (Molecular Devices). Signals were sampled at 10 kHz and filtered at 2 kHz. Data were stored on a PC for offline analysis after digitization using an A/D converter (Digidata 1440A, Molecular Devices).

### Image analysis

To measure the stability of microdomains, colocalization analysis was performed by selecting a region of interest (ROI) and measuring YFP/CFP in each pixel of that ROI utilizing ImageJ (Figure S5A, B). Pixel intensity correlation analysis in time-lapsed images was performed by generating 2D intensity histograms and calculating a pixel-to-pixel Pearson’s r between sequential images, using the Coloc2 Fiji subroutine (Figure S5C). Pseudo-colored images of <u>sCLM</u> (Figure 2A, C, D) were generated by selecting the ROIs, measuring YFP/CFP in each pixel of that ROI and using the Pseudo-color Image Look-Up Tables (LUTs) Fiji subroutine. The remaining pseudo-colored images of <u>sCLM</u> (Figure 2F and S4E) were generated by reconstructing a flattened (2D) image using the Cl^−^ insensitive CFP intensity to weight the contribution of each pixel’s YFP/CFP ratio in a 3D image stack. Pseudo-color images of FLIM (Figure 3E and S4B) were generated by calibrating MEQ lifetime utilizing the Stern-Volmer relationship and calculating [Cl^−^]_i_ (Figure S4B) (Verkman et al., 1989). FLIM experiments included the brighter membrane impermeable dye Alexa Fluor-594 as well as MEQ in the recording pipette solution. The Alexa Fluor-594 (Cl^−^-insensitive) signal was used to define the intracellular pixels. [Cl^−^]_i_ was then calculated from the fluorescence lifetime of MEQ in those pixels. A 2D image was created from a 3D image stack by weighting the MEQ-derived [Cl^−^]_i_ values by the relative intensity of Alexa Fluor-594. The Alexa Fluor-594 signal intensity was used to determine the brightness of each pixel that was pseudo-colored based on the calculated [Cl^−^]_i_. In pixels where MEQ signal is indistinguishable from background noise, no pseudo-color is assigned, and the pixel is colored using a grey-scale Alexa Fluor-594 signal. The greyscale then demonstrates the dendritic structure in areas where MEQ signal is inadequate. For noise analysis, a representative 30-minute time series (180 frames) of acutely sliced CLM-expressing CA1 pyramidal cells imaged in the presence of TTX is compared to the noise acquired from a similar series acquired with the shutter closed (Figure 2H). MATLAB is used to perform a line-scan to represent each neuron as a vector of intensity values and to compute subsequent Fast Fourier Transforms (FFTs) of CLM YFP intensity and the corresponding channel closed-shutter noise intensity (see detailed noise analysis below). As the power spectrum of more timeframe images are averaged, the power of noise from a closed shutter will converge to that of white noise. Power obtained from variations due to persistent structural differences such as microdomains will not converge to a white noise power spectrum, converging instead to a greater power difference from white noise than shutter noise (Figure 2H).

### Detailed noise analysis

To test whether the observed spatial variance in [Cl^−^]_i_ could arise from noise, we compared the averaged power spectra of the spatial frequencies of [Cl^−^]_i_ reporter CLM intensity values in the neurons to the power spectrum of closed-shutter noise. Noise analysis was performed with customized scripts in MATLAB utilizing Fourier transform and image analysis functions therein. Of the two fluorescent components of CLM, Cl^−^-sensitive YFP was chosen and compared to the noise detected with the laser shutter closed over the same number of timeframes for each neuron. Dendrites or soma were chosen based on persistent signal strength and linearized by line scan. The YFP signal of each frame was co-registered to the first frame and applied to both the YFP and closed-shutter noise image of that frame. Spatial Fast Fourier Transforms (FFTs) were computed on these intensity signals to obtain the power (amplitude squared) of the observed YFP and closed-shutter noise values at a spectrum of wavelengths from 0.794 μm (exclusive) to 25.4 μ m (inclusive), bound by the Nyquist wavelength and the distance equivalent to the largest exponent of base 2 that did not exceed the length of the smallest vector in pixels, respectively. The closed shutter noise power spectra are rescaled by the ratio of area under the curves between the mean power spectrum of the neuronal signal and shutter noise. The white noise power spectrum was generated from averaging the power spectra of 72,000 vectors created from 400 random shuffles of each of 180 timeframes. Defining *X*_1…180_ as the vectors at 180 different timeframes of observed intensity values for noise from a closed shutter or a signal from a neuron, we obtain 400 randomly shuffled versions for each 180 timeframe vectors *X*_1…180_ assigned to *A*_1…72,000_. By computing the Fast Fourier Transform (FFT) on each of the vectors *A*_1…72,000_ and averaging the square of each result, we obtain the respective white noise power spectrum *W* for either a neuronal signal or closed shutter noise defined as:

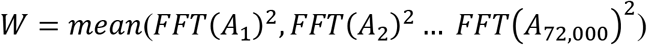

The absolute area between the cumulative average of either the power spectra of the YFP neuronal signal or the closed-shutter noise and their respective white noise power spectrum is calculated at each timeframe. The resulting curves are normalized and plotted as a function of frame number such that the number of frames included in the average power spectrum is equal to the frame number. Comparing the behavior of the averaged power spectra of signal intensity versus closed-shutter noise as a function of the number of frames averaged allows a direct comparison of heterogeneity. Heterogeneity that is random and transient (i.e. noise) will average out with increasing number of frames, but heterogeneity that is persistent will not.

### Quantification and statistical analysis

Imaging data were analyzed with ImageJ (http://imagej.nih.gov/ii\\j/) or MATLAB. Details of the analysis are included in “image analysis” section of STAR Methods. Statistical analysis of the data was done in GraphPad Prism (GraphPad Prism 8). Statistics were assessed with two-tailed unpaired Student’s t tests when comparing two groups. The number of data points (n) and the statistical significance (p value) are stated in figure legends.

## Supplemental Information

**Supplemental Table 1:**
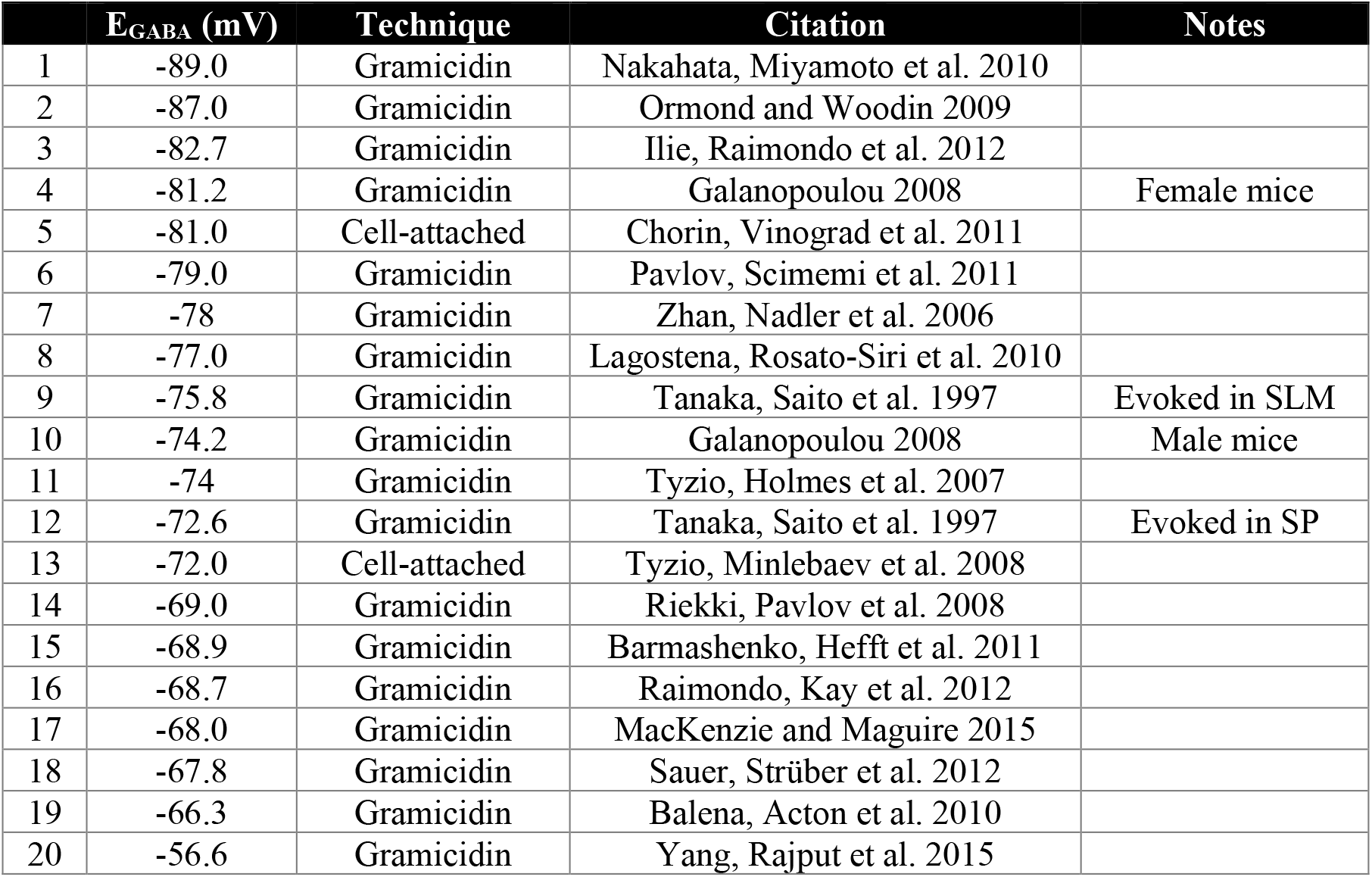
Reported range of E_GABA_ in separated studies. 20 group means of E_GABA_ measurements using gramicidin and cell-attached techniques in age-matched CA1 pyramidal cells. SP: Stratum pyramidale, SLM: stratum lacunosum-moleculare.

**Supplemental Figure 1:**
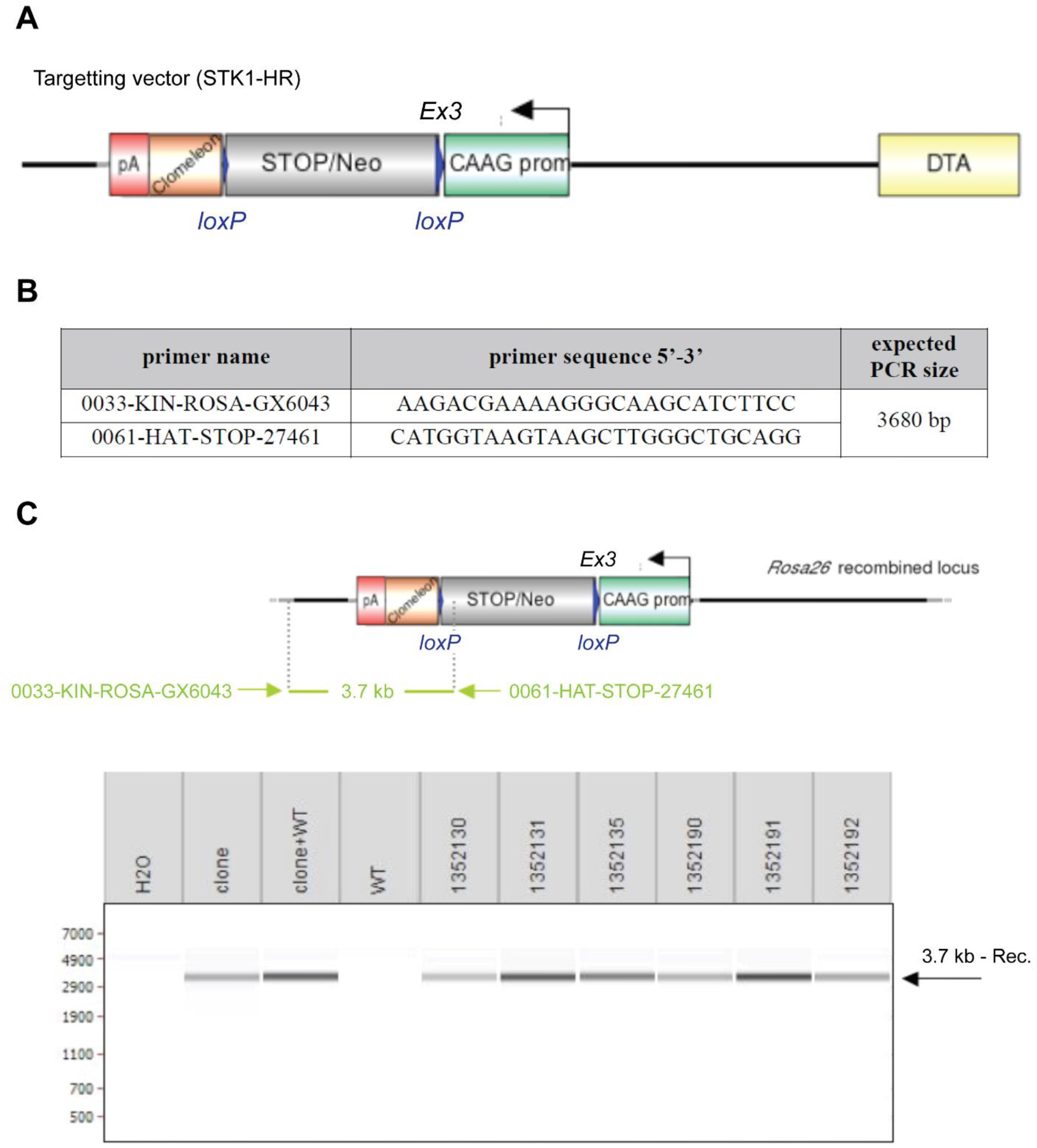
Construction of the *sCLM* targeting vector and Generation of chimeric mice. **A)** Schematic representation of Rosa26 Knock-in targeting vector. Diagram is not depicted to scale. LoxP sites are represented by blue triangles. The sCLM transgene is depicted as a brown box and the polyA as a red box. The combined STOP-neomycin selection cassette is indicated as a grey box. The ubiquitous CAAG promoter is represented by a green box. The DTA negative selection cassette is represented by a yellow box. **B)** primer set for the detection of the non-excised recombined conditional Knock-in allele. **c.** The genotype of the pups derived from the F1 breeding with Cre deleter mice were tested by PCR enabling the detection of the neomycin cassette. PCR using recombined ES cell clones as template, alone or in the presence of 10 ng C57BL/6 genomic DNA (clone; clone+WT) served as positive controls. PCR with wild-type C57BL/6 genomic DNA (WT) and without DNA (H2O) as template served as negative controls. PCR picture was obtained after loading of PCR reactions on LabChip® system from Caliper LifeSciences.

**Supplemental Figure 2:**
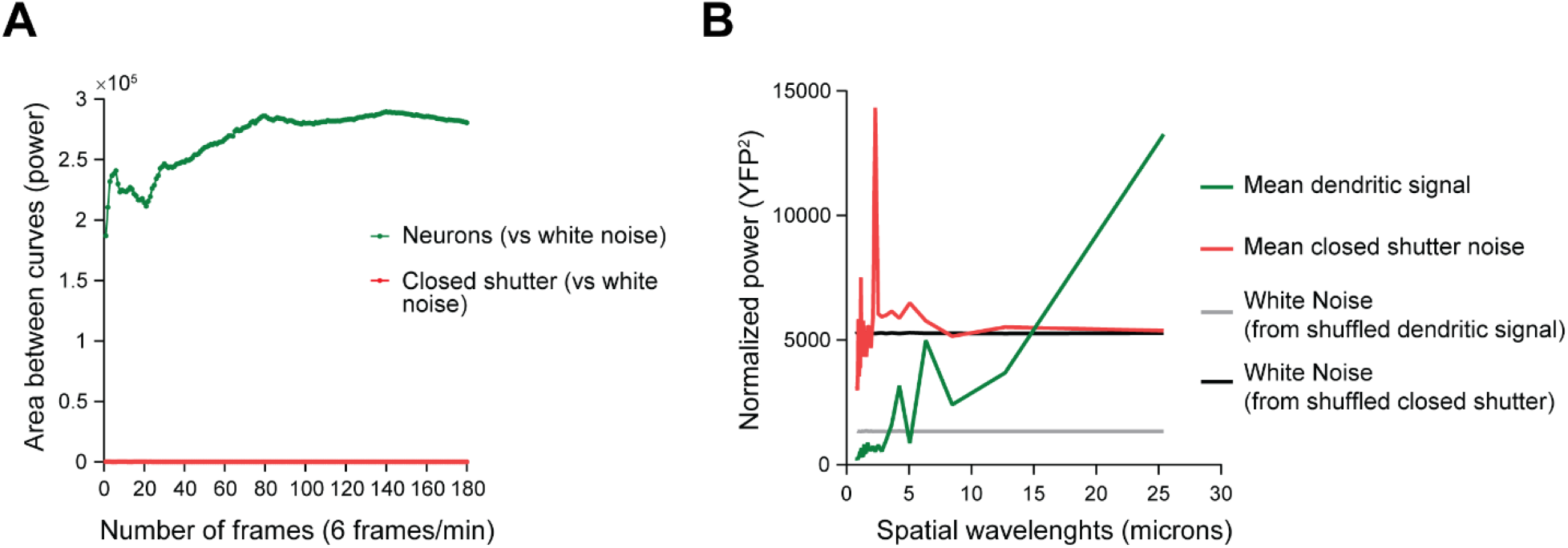
Signal vs noise analysis. **A)** From 180 Clomeleon timeframes, the absolute area between the mean power spectra and that of white noise for both the neuronal YFP fluorescence signal and closed shutter noise is plotted against the number of timeframes included in the mean (green and red, respectively). These are the raw curves which are then normalized in panel B. **B)** The mean power spectrum of closed shutter noise is normalized by a ratio to have total power equivalent to that of the mean power spectrum of the YFP dendritic signal (red and green, respectively). A white noise power spectrum is generated for either normalized shutter noise or dendritic signal (black and gray, respectively).

**Supplemental Figure 3:**
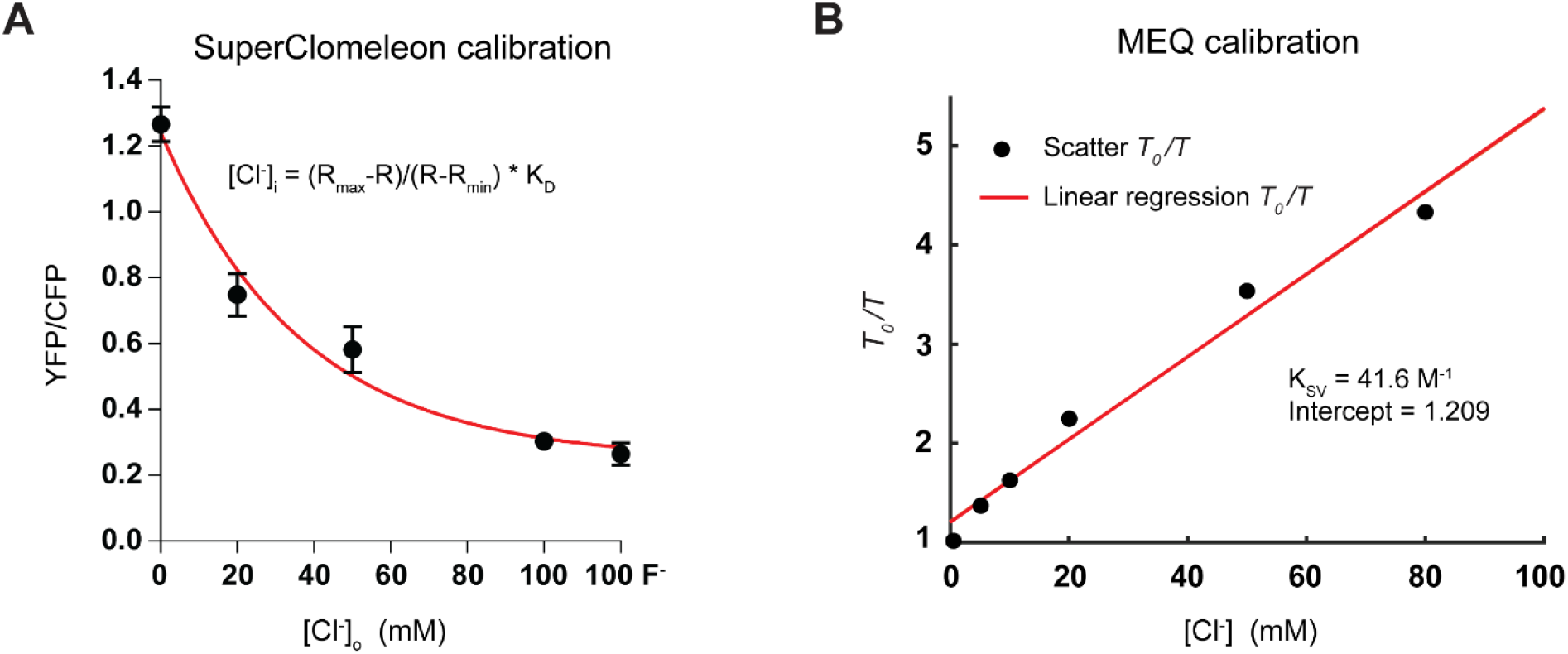
Calibration of SuperClomeleon and MEQ. **A)** SuperClomeleon was calibrated in organotypic slice cultures at different concentrations of Cl^−^ and in the presence of 50 μm nigericin and 100 μm tributyltin. With single-cell resolution, we calculated R_max_, R_min_ and K_D_ as described in Methods section. Only neurons that responded to different Cl^−^ concentrations in extracellular media ([Cl^−^]_o_) were selected for final calculations. The range of values for R_max_ measured by two-photon microscopy in different calibration experiments was between 1.240 and 1.777. R_min_ ranged between 0.289 and 0.466. K_D_ was between 22.31 and 22.99 mM. For DualView imaging, R_max_, R_min_ were 1.640, 0.984 respectively. **B)** MEQ was calibrated using Stern-Volmer equation. MEQ was dissolved in the same solution as the internal pipette solution (K-MeSO_4_-based) with different concentrations of Cl^−^.

**Supplemental Figure 4:**
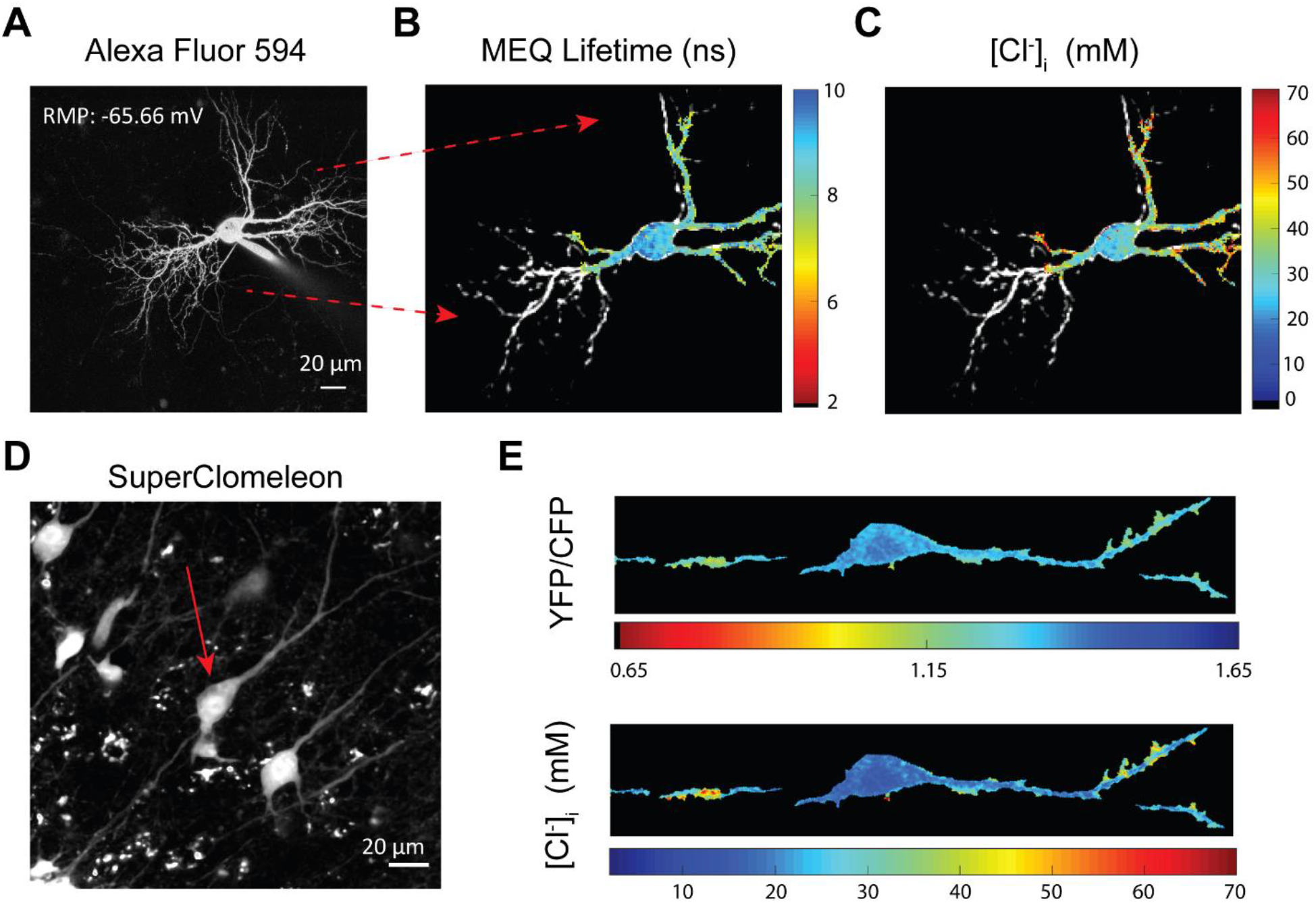
Extra examples of [Cl^−^]i two photon and lifetime imaging. **A)** Two-photon imaging of a pyramidal cell loaded with Alexa Fluor-594. **B)** Same neuron in A, was also loaded with MEQ to measure lifetime. **C**) [Cl^−^]_i_ was calculated based on MEQ calibration shown in Figure S2. **D)** Two-photon imaging of an interneuron expressing SuperClomeleon. **e)** The calculated YFP/CFP ratio and [Cl^−^]_i_.

**Supplemental Figure 5:**
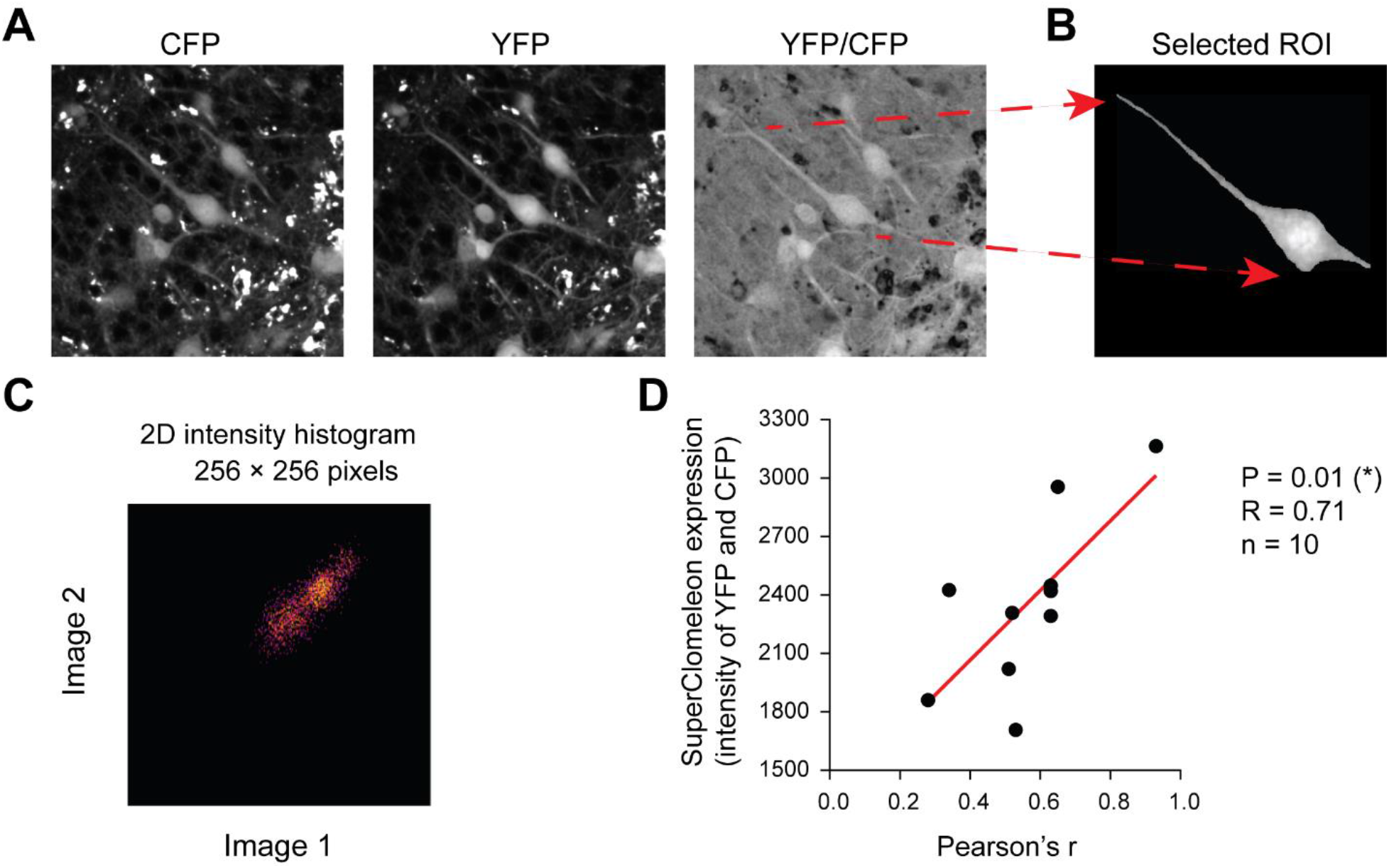
Measuring stability of microdomains. **A, B)** Colocalization analysis was performed by selecting a region of interest (ROI) and measuring YFP/CFP in each pixel of that ROI. **C)** Pixel intensity correlation analysis in time-lapsed images was performed by generating 2D intensity histograms and calculating Pearson’s r, using Coloc2 Fiji’s plugin. **D)** The observed variability in Pearson’s r in different experiments was significantly correlated with expression levels of SuperClomeleon.

## Notes

### Competing Interest Statement

The authors have declared no competing interest.

